# Learning to Choose: Behavioral Dynamics Underlying the Initial Acquisition of Decision Making

**DOI:** 10.1101/2024.02.28.582581

**Authors:** Samantha R. White, Michael W. Preston, Kyra Swanson, Mark Laubach

## Abstract

Current theories of decision making propose that decisions arise through competition between choice options. Computational models of the decision process estimate how quickly information about choice options is integrated and how much information is needed to trigger a choice. Experiments using this approach typically report data from well-trained participants. As such, we do not know how the decision process evolves as a decision-making task is learned for the first time. To address this gap, we used a behavioral design separating learning the value of choice options from learning to make choices. We trained male rats to respond to single visual stimuli with different reward values. Then, we trained them to make choices between pairs of stimuli. Initially, the rats responded more slowly when presented with choices. However, as they gained experience in making choices, this slowing reduced. Response slowing on choice trials persisted throughout the testing period. We found that it was specifically associated with increased exponential variability when the rats chose the higher value stimulus. Additionally, our analysis using drift diffusion modeling revealed that the rats required less information to make choices over time. Surprisingly, we observed reductions in the decision threshold after just a single session of choice learning. These findings provide new insights into the learning process of decision-making tasks. They suggest that the value of choice options and the ability to make choices are learned separately, and that experience plays a crucial role in improving decision-making performance.

## INTRODUCTION

Researchers studying decision-making in animals utilize behavioral tasks that often require extensive training to reduce behavioral variability. As such, neuroscientific studies of decision-making have traditionally recorded neural activity from subjects with abundant experience in making choices (Carandini and Churchland, 2013). For instance, studies employing two-alternative forced-choice (2AFC) paradigms train animals to discriminate between stimulus pairs and record brain activity after animals reach high levels of behavioral performance (Zoccolan et al., 2009; Busse et al., 2011; Brunton et al., 2013; Reinagel, 2013; Erlich et al., 2015; Hanks et al., 2015; Burgess et al., 2017; Kurylo et al., 2020; Masis et al., 2023). These studies provide valuable insights into neural and computational mechanisms underlying decision making, but have not addressed how the training process may influence decision making strategies or neural activity.

Neural and computational models of decision making assume an internal comparative process when participants are faced with multiple options. In studies using 2AFC task designs, participants are trained to learn the value of simultaneously presented stimuli. Choosing one while forgoing the other may ensure the adoption of a comparative strategy. Few, if any, studies have addressed how decision making tasks are learned if participants learn the reward values of task stimuli prior to making choices between the stimuli. We wondered whether the training procedures used for 2AFC tasks might shape the learning process and influence decision making strategies.

Studies reviewed by Kacelnik et al. (2011) highlight this concern. Their research with European starlings revealed that, in nature, animals typically encounter food options sequentially, not simultaneously as presented in many lab experiments. To better mimic natural foraging in the laboratory, they used a unique design (Shapiro et al., 2008). During training, visual stimuli were presented individually (sequentially) on some trials and together (simultaneously) on other trials. Kacelnik and colleagues found that choices between the stimuli were predicted by the animals’ latencies to respond to the single offers of the stimuli. They further found no differences between the time taken to make a choice between pairs of stimulus and the time taken to respond to offers of just one of the stimuli. Their studies suggest a lack of deliberation, or slowing of response times, on choice trials. This finding was replicated in subsequent studies involving rodents (Ojeda et al., 2018; Ajuwon et al., 2023). Kacelnik et al. (2011) argue that the act of deliberation, observed in lab settings using two-alternative forced choice designs, might be an artifact of the training process. They further suggest that Drift Diffusion Models (Ratcliff, 1978), which rely on competition between stimulus options, may not be suitable for understanding decision-making. These findings necessitate reevaluating current interpretations of decision-making research, which often rely on the assumption that deliberation slows down choices and that Drift Diffusion models adequately capture the underlying neural processes.

Building on the work of Kacelnik et al. (2011), we designed a behavioral task to isolate learning the values of individual stimuli from learning to make choices between stimuli (Fig 1). Using an open-source two-alternative forced-choice task (Swanson et al., 2021), we first trained rats to learn the reward values of the stimuli. Then, we introduced choice trials with pairs of stimuli and measured how the rats responded when making choices for the first time. Contrary to findings summarized in Kacelnik et al. (2011), responses were initially slower in the first session with choice trials compared to the single-option trials. We continued testing over several sessions to assess the effect of experience on decision-making. Interestingly, even as the rats became faster overall, the initial slowing down during choice trials persisted, suggesting a separate process for decision-making beyond reward learning.

**Figure 1:**
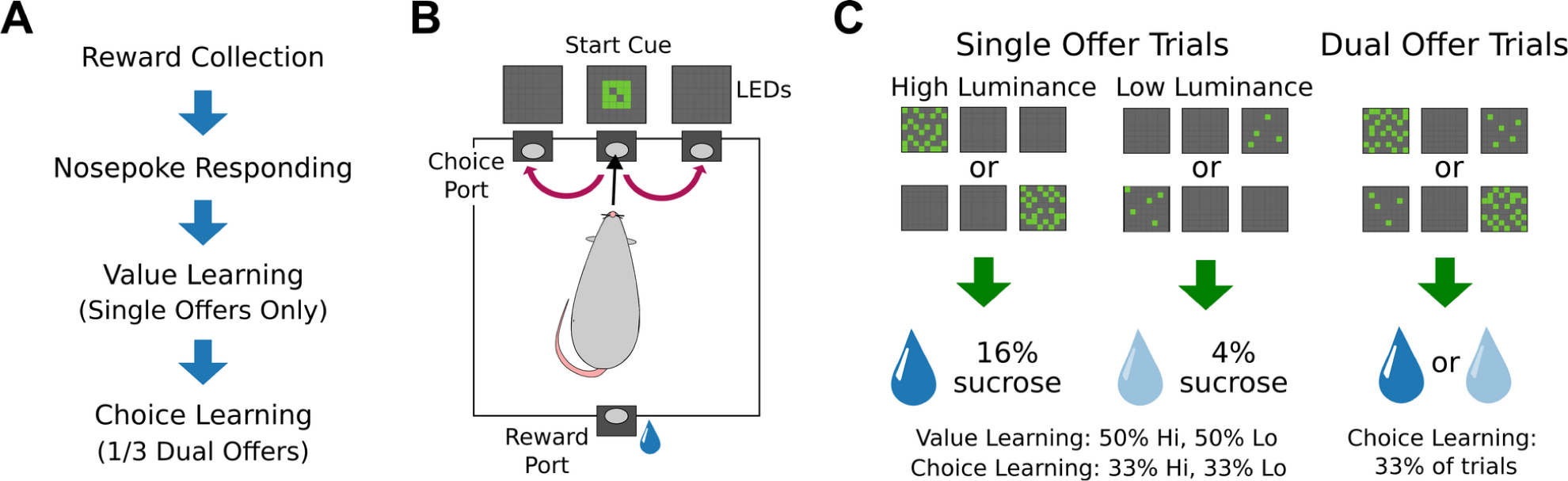
Task training, design, and trial types. A) Rats went through a series of intermediate training steps to learn the task protocol, including how to collect the liquid sucrose reward and nosepoke to gain access to said reward, and learn cue values (“value learning”) for several sessions before experiencing “choice learning”. B) Animals responded to visual cues on one side of the operant chamber and crossed the chamber to consume liquid sucrose at the reward spout. We measured response latency as the time elapsed between trial initiation (center poke) to nosepoking the port with the visual stimulus (red arrows). C) For the value learning phase, rats only had access to single-offer trials of either high or low luminance stimuli. Upon entering the central port on these trials, one cue would appear above either the left or right nosepoke ports, randomized by location and value. Responding to the high luminance stimulus led to access to 16% liquid sucrose at the reward port. Responding to the low luminance stimulus led to access to 4% liquid sucrose. During testing in the choice learning phase, 2/3 of the trials consisted of these single-offer trials. For the remaining 1/3 of trials, rats were exposed to dual-offer trials with both high and low luminance cues displayed. Single and dual offers were randomly interleaved.

To analyze the decision-making process during initial choice learning, we employed two computational models. First, we used ExGauss models to assess the impact of learning on response time distribution (Hohle, 1965; Luce, 1986). This revealed how learning influenced the overall speed and variability of choices. Second, we employed drift diffusion models to explore if early learning modified parameters within this common neuroscience framework. Our analysis showed that initial learning primarily influenced the decision threshold parameter, not others, within the drift diffusion model. Interestingly, these changes in threshold positively correlated with the exponential variability in response times, which has been linked to noise in the decision process (Hohle, 1965). These findings suggest that early learning may lead to needing less information to make a choice and potentially reducing noise in the decision process.

## RESULTS

### Rats deliberate when making choices for the first time

We sought to investigate whether training rats with single-offer trials and then exposing them to dual-offer trials with known stimuli would produce behavioral changes associated with deliberation. After training for multiple sessions with single-offer trials, rats show marked differences in behavior on dual-offer trials. In a one hour session, rats on average completed 301+/-74 trials. Rats chose the high-value stimuli (72%+/-6%) more often than the low-value stimuli (28%+/-6%) when both options were presented during the initial test session (t(24)=-16.16, p<0.001; paired t-test) (Fig 2A). We also found that rats show an increased median response latency for trials with dual offers compared (625ms) to trials with single offers (521ms) (F(1,24)=37.36, p<0.001; rmANOVA) and for trials with low-value stimuli (597ms) compared to trials with high-value stimuli (527ms) (F(1,24)=28.31, p<0.001; rmANOVA) (Fig 2B). Further, when broken out by value, the difference between dual- and single-offer median latencies was significant for high-value trials (mean difference: 0.09, p<0.001; paired permutation test) and low-value trials (mean difference: 0.12, p<0.001; paired permutation test). When looking at the response distributions, we found that trials with single and dual offers generally had non-overlapping exponential tail distributions (Fig 2C). Finally, we assessed within-session effects on latencies and found there was a general effect of slowing over the one hour session (1H: weighted *R*^2^=0.25, F(1,78)=4.397 p<0.001; 1L weighted *R*^2^=0.42, F(1,72)=4.082 p<0.001: 2H: weighted *R*^2^=0.17, F(1,57)=4.563 p<0.001; Robust M-Regression) (Fig 2D).

**Figure 2:**
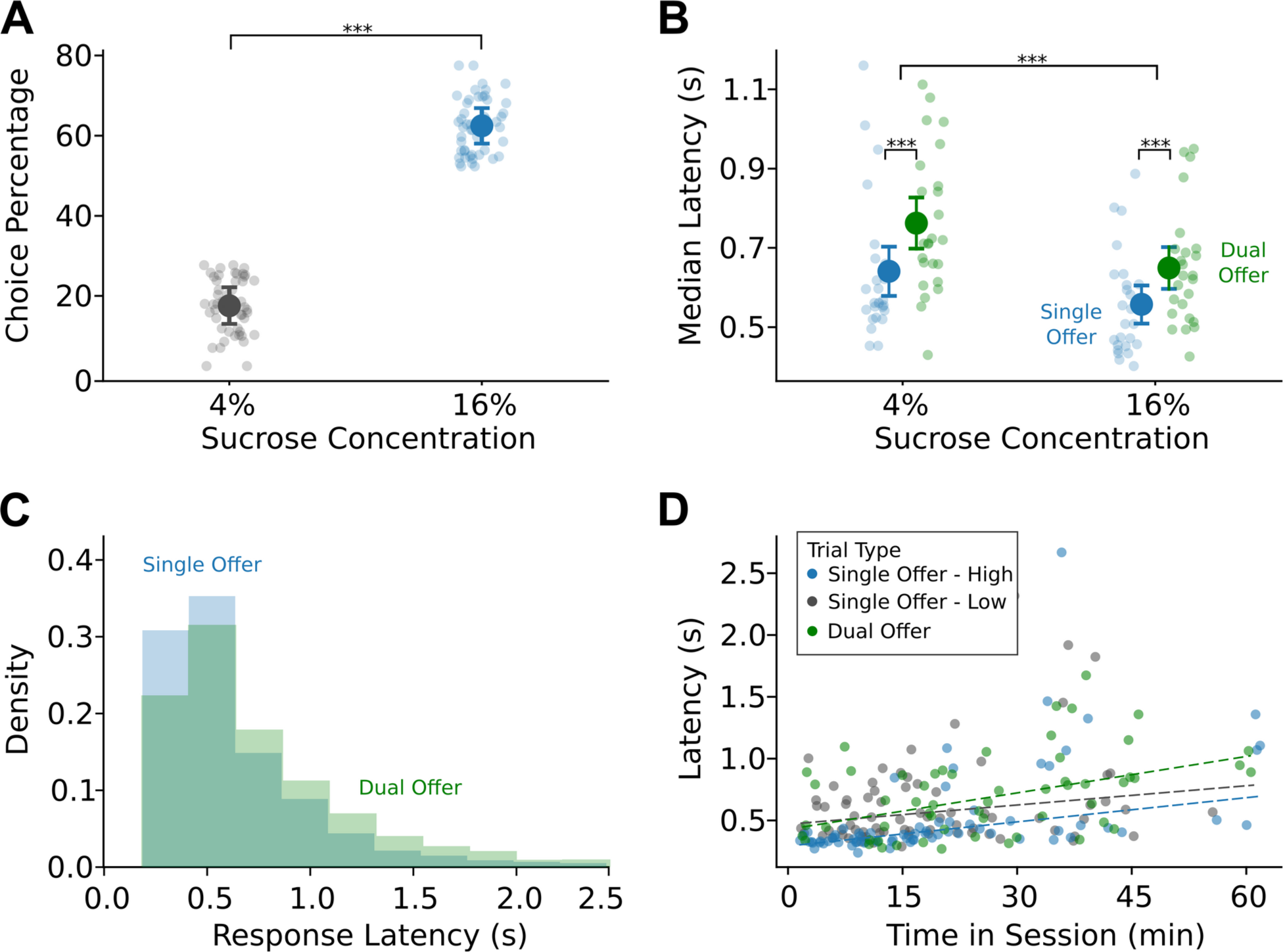
Rats deliberate when making choices for the first time. A) On dual-offer trials, rats (N=25) selected the high luminance, high-value cue approximately 72% of the time overall, suggesting these subjects prefer the high-value reward over the low-value option. B) Rats showed overall increased latencies for low-value offers compared to high-value offers. Importantly, rats showed increased latencies for dual-offer trials compared to single-offer trials regardless of the chosen value. C) Raw latency distributions for trials with single and dual offers showed a non-overlapping portion in the tail ends of the distribution. D) Response latencies generally increased over the session across trial types and values (data from one exemplar rat shown).

### Deliberation is persistent over the period of early choice learning

Given the robust effect that we found with respect to latency differences from trials with single and dual offers, we wanted to examine how stable the deliberation effect was with repeated testing. Fifteen rats were tested over five sessions of choice learning (Fig 3). Given the overall differences in latency between high and low-value stimuli (F(1,14)=87.06, p<0.001), we broke out trials by value and used Bonferroni corrections for tests of significance. In the case of high-value trials, we found an overall effect of session number on median latencies (F(1,14)=6.953, p<0.001; rmANOVA) and an overall effect of trial type (F(1,14)=34.607, p<0.001; rmANOVA).

**Figure 3:**
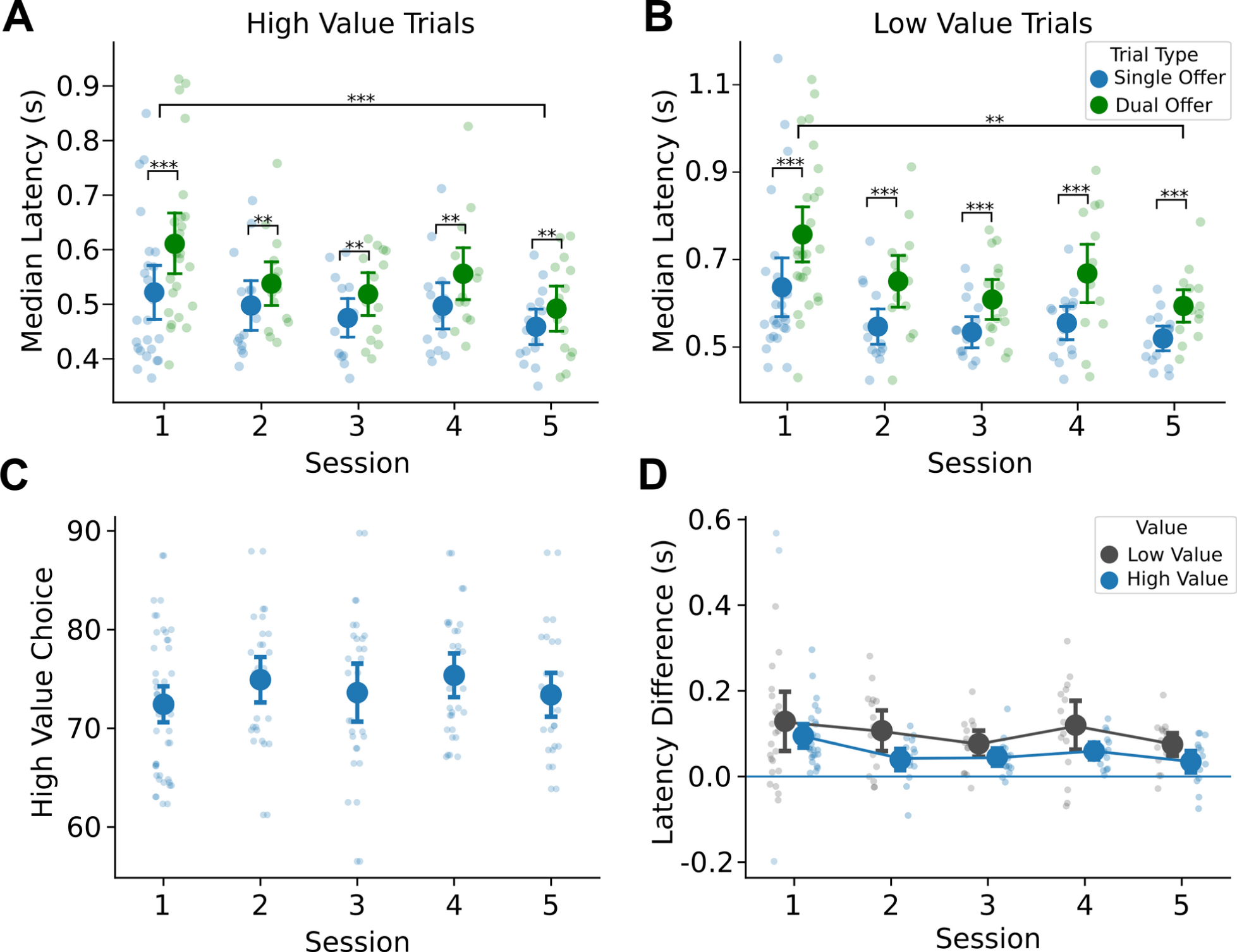
Rats reduce their response latencies but maintain deliberation with experience in making choices. A) Median response latencies for high-value trials from 15 rats showed reduction with experience while maintaining an increase on dual-offer compared to single-offer trials. B) Median response latencies for low-value trials were overall greater than high-value trials and showed reduction with experience while maintaining an increase on dual-offer compared to single-offer trials. C) Rats did not change their proportion of high-value choices on dual-offer trials with more experience. D) The magnitude of difference in single- and dual-offer median latencies reduced with experience, especially from the first to second session, however, subjects maintained an elevated latency for dual-offer trials over the course of choice learning.

To further investigate the differences in latencies for trials with single and dual offers, we used paired permutation tests with Bonferroni correction given this test is robust for relatively small sample sizes. We found that the difference in median latencies from trials with single and dual offers persisted over the 5 sessions (see Fig 3A; paired permutation tests). For low-value trials, we found similar results; overall effect of session number on median latencies (F(1,14)=4.593, p=0.00168; rmANOVA), an overall effect of trial type (F(1,14)=57.750, p<0.001; rmANOVA), and the difference in median latencies from trials with single and dual offers persisted over the 5 sessions (Fig 3B; paired permutation tests). These changes in median response latency occurred in the absence of an effect on high-value choice percentage (F(1,14)=2.339, p=0.0585; rmANOVA) (Fig 3C).

To emphasize the effect that dual-offer trials has on response latencies, we analyzed the median difference in trial types by value and confirmed that these latency differences persist over 5 sessions, however, there was a greater latency difference in the first session compared to the subsequent sessions (High value: F(1,14)=3.171, p= 0.0203; Low value: F(1,14)=3.503, p=0.0127; rmANOVA) (Fig 3D).

### Computational analyses of response latencies and decision dynamics

Two computational analyses were used to understand how early choice learning influenced the animals’ response latencies and the decision process. The first analysis examined the skewed nature of the latency distributions (Fig 2C) using ExGauss modeling (Heathcote et al., 1991). ExGauss models tease apart the latency distributions into Gaussian components (mu parameter) that account for the peaks in the distributions and exponential components (tau parameter) that account for the positively skewed tails in the distributions (Fig 4A). These measures have been interpreted, respectively, as reflecting sensorimotor processing and variability in the decision process (Hohle, 1965; Luce, 1986). The second analysis used hierarchical Bayesian drift diffusion modeling (HDDM; Wiecki et al., 2013) and generalized drift diffusion modeling (PyDDM; Shinn et al., 2020). This approach allowed us to measure how early choice learning affected the rate of evidence accumulation prior to a choice (drift rate) and the amount of information needed to make a choice (threshold), as well as the time required for stimulus and motor processing (non-decision time) (Fig 4B). By comparing parameters from the two types of models (ExGauss and DDMs), we were able to assess how early choice learning affected the statistical properties of the response latencies across sessions and different types of trials (i.e., responses to stimuli with high and low reward values, responses on trials with single and dual offers) and how the statistical properties of the response latencies varied in relation to the three main decision variables (drift rate, threshold, and non-decision time).

**Figure 4:**
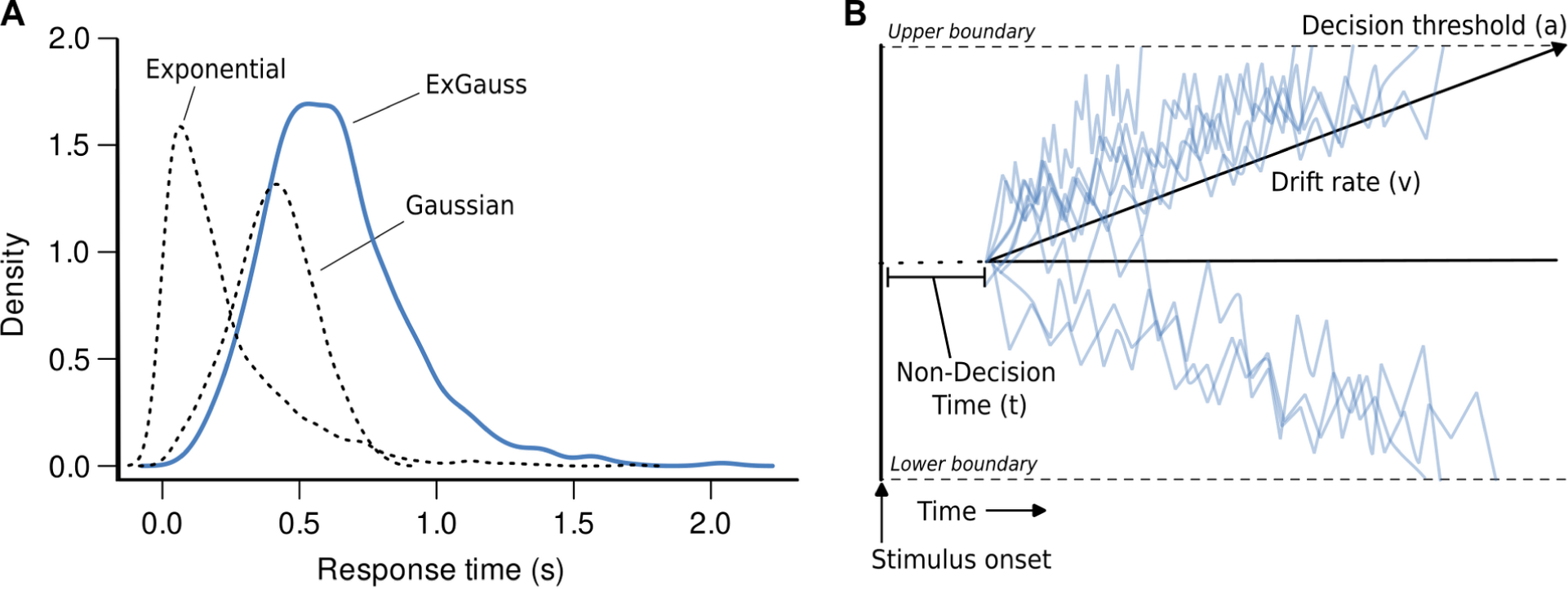
ExGauss and Drift Diffusion Modeling. A) ExGauss models estimate response time distributions as mixtures of Gaussian and exponential distributions. The ExGauss distribution is the sum of the Gaussian and exponential components. In the example shown here, parameters from one of the rats was used to simulate full distributions for each component. The Gaussian component accounts for the peak in the response time distribution. The exponential component accounts for the long positive tail in the distribution. B) Drift Diffusion Models, or DDMs, estimate three parameters of the decision process based on random fluctuations of evidence for the response options over the time from the onset of the stimuli to the time of choice. The drift rate reflects the slope of the accumulation of evidence towards the threshold, when a choice is triggered. Non-decision time accounts for stimulus integration and sensorimotor processing.

### ExGauss modeling of response time distributions

We fit ExGauss models to the response latency distribution of each rat for each learning session and each type of trial. We found that the rats showed more variable latencies when choosing the high-value stimulus compared to when they were forced to respond to the stimulus (Fig 5A,C). By contrast, they responded more slowly, with equal variability, when choosing the low-value stimulus compared to forced responses to that stimulus (Fig 5B,D). For high-value trials, we found an overall effect of session number on the Gaussian component (F(1,14)=4.170, p=0.00328; rmANOVA) (Fig 5A), but no overall effect of trial type (F(1,14)=2.915, p=0.09016; rmANOVA). In the case of low-value trials, however, we found an overall effect of both sessions (F(1,14)=5.076, p<0.001; rmANOVA) and trial type (F(1,14)=18.438, p<0.001; rmANOVA) (Fig 5B). Paired permutation tests further reveal a persistent difference between trials with single and dual offers over the first four sessions.

**Figure 5:**
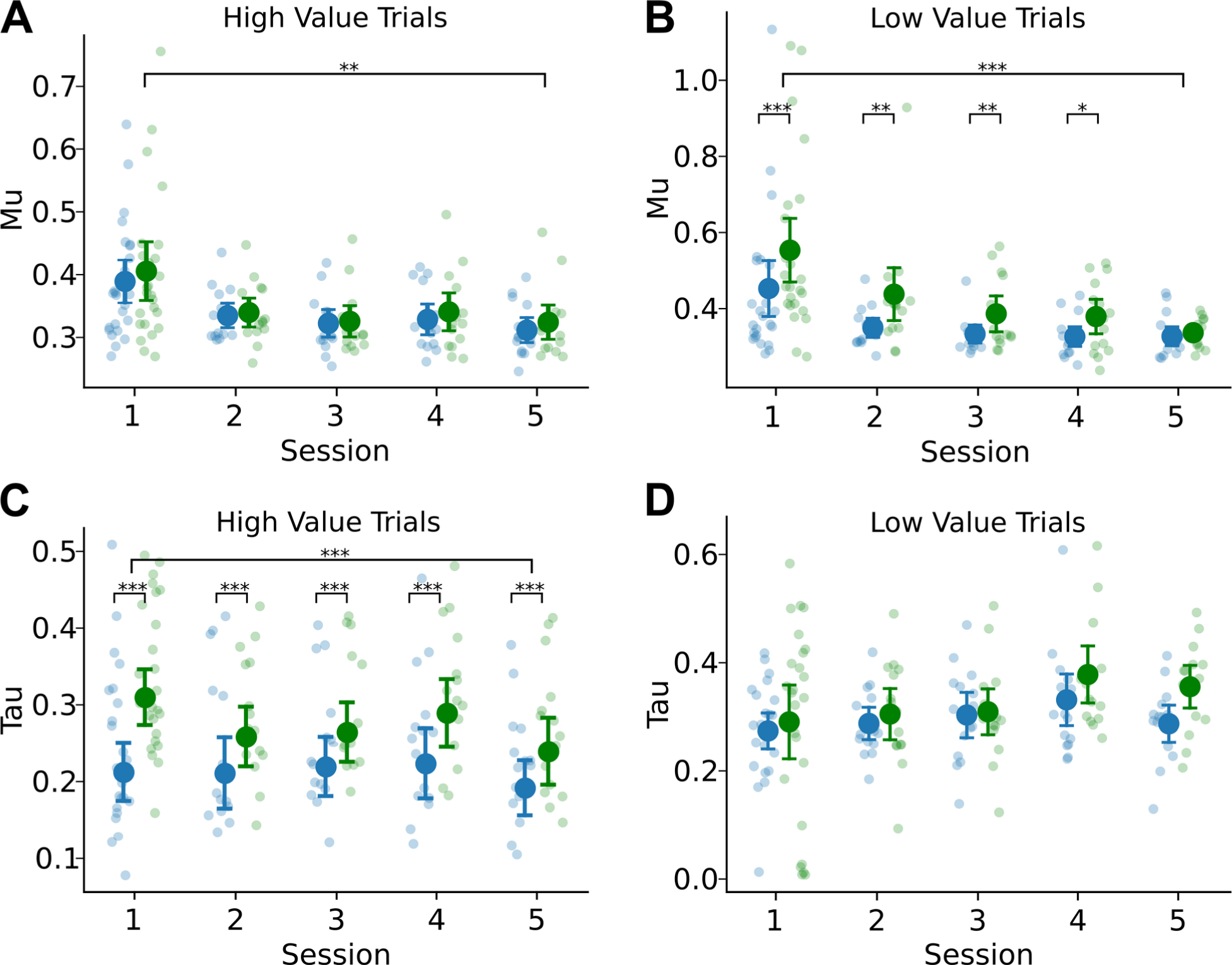
ExGauss modeling reveals differences in response latency distributions for dual- and single-offer trials. A) The ExGauss parameter Mu represents the mean of the Gaussian component, a measure of sensorimotor integration. An increase in this parameter indicates an overall shift in the distribution to the right. There was no difference for single and dual trial fits of the Mu parameter for high-value trials, however there was an overall reduction in the Mu parameter over all the sessions. B) In the case of low-value trials, there was an overall increase of the Mu parameter on dual-offer trials and a decrease in this parameter over all the sessions. C) The ExGauss parameter Tau represents the mean of the exponential component, a measure of variability in the decision process. An increase in this parameter indicates an overall lengthening of the distribution tail to the right. There was an overall increase of the Tau parameter on dual-offer trials for the high-value stimulus, and a decrease in this parameter over all the sessions. D) In the case of low-value trials, there was no difference for single and dual trial fits of the Tau parameter and no change over the sessions.

When examining the response latency variability in the exponential tail, we found the opposite effect with respect to value. For high-value trials, we found an overall effect of session (F(1,14)=4.254, p=0.00287; rmANOVA) and an overall effect of trial type (F(1,14)=37.559, p<0.001; rmANOVA) (Fig 5C). Paired permutation tests further reveal a persistent difference between trials with single and dual offers over the five sessions. In the case of low-value trials, there was no effect of session (F(1,14)=1.651, p=0.165; rmANOVA) nor trial type (F(1,14)=2.553, p=0.113; rmANOVA) on the exponential variability (Fig 5D).

### Drift diffusion modeling of decision dynamics

To further explore how decision-making processes change over sessions, we fit two versions of drift diffusion models (HDDM and PyDDM) using the animals’ choices and response latencies from dual-offer trials (Fig 6). A common finding across models was that initial choice learning reduced the decision threshold. Importantly, this effect was observed after a single session of choice learning (Fig 6B).

**Figure 6:**
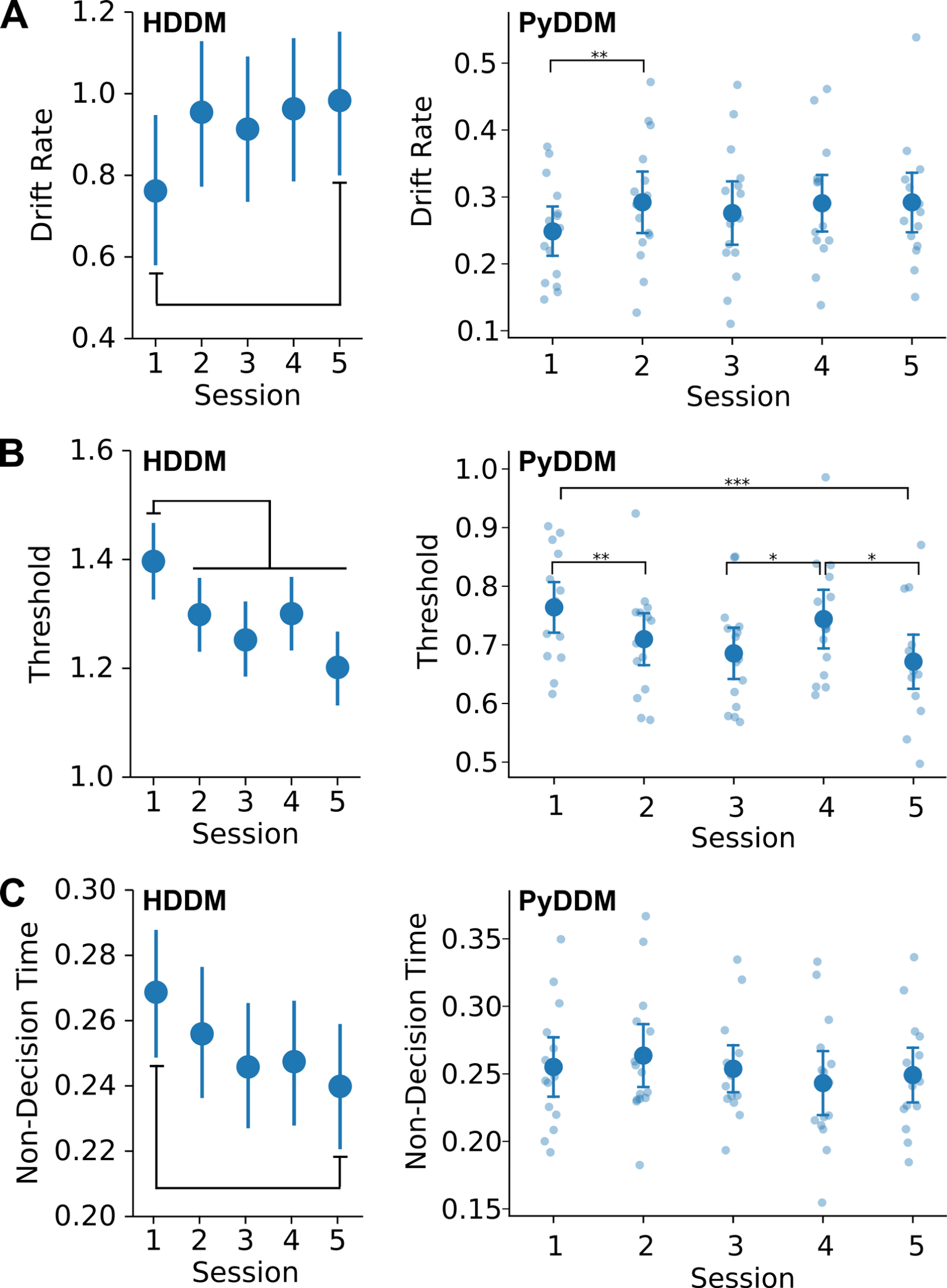
Rats required less evidence to make choices during early learning. A) Drift rate increased after the first session of choice learning. B) Threshold reduced over the period of choice learning, and importantly was reduced after a single test session. C) Non-decision time based on HDDM was higher in the first test session compared to the fifth session. No effects of learning on non-decision time were found using PyDDM. Please note that the two kinds of DDMs report the decision parameters using different units.

The left plots in each panel in Fig 6 show Bayesian estimates of the mean and 95% confidence intervals for the DDM parameters from analysis using HDDM. Differences are noted based on comparisons of the posterior distributions for each parameter, comparing the first session against the rest. Drift rate was lower in the first learning session compared to the fifth session (P(1>5): 0.0456, Fig 6A, left). Threshold was higher in the first learning session compared to all other sessions (e.g., P(1<2): 0.0204, Fig 6B, left). Non-decision time was longer in the first learning session compared to the fifth session (P(1<5): 0.0172, Fig 6C, left).

Fig 6 also shows results from the PyDDM models. We found no overall effect of session number on the drift (Fig 5B; F(1,14)=1.979, p=0.11; rmANOVA) or non-decision time (Fig 5D; F(1,14)=1.382, p=0.252; rmANOVA) parameters. We did, however, find a significant increase in drift from the first to second session before the drift stabilizes for the remaining sessions (paired mean difference: 0.044, p=0.0078). We found an overall effect of the boundary separation parameter (Fig 5C; F(1,14)=4.316, p=0.0041; rmANOVA), with differences from session to session (see Fig 5C).

### Relationships between measures of choice behavior and the drift diffusion models

We used repeated-measures correlation to assess how the behavioral and computational measures reported above related to each other. For this analysis, we used the parameters from the PyDDM models and related them to the animals’ preferences for the higher value stimulus and measures of their response times based on ExGauss modeling. The strongest correlation across measures was between the drift rate and the percent of trials in which the rats chose the higher value stimulus (r=0.9222, df=69, p<0.001, CI95%: 0.88–0.95) (Fig 7A).

**Figure 7:**
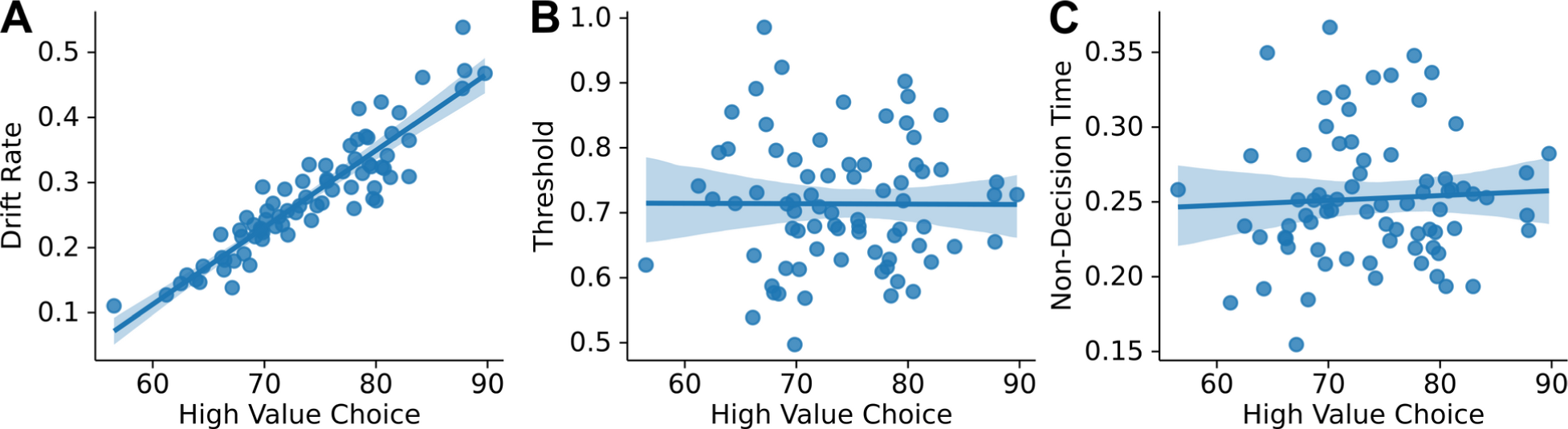
Drift rate is driven by choice preference. Scatterplots are shown for the high-value choice percentage across animals and sessions versus the three parameters from the PyDDM models. Regression lines were fit and show 95% confidence intervals. Correlational values were calculated using repeated-measures correlation to control for effects of training session. A) There was a strong positive relationship between choice preference and drift rate. B,C) There was no clear relationship between choice preference and threshold or non-decision time.

Threshold and non-decision time did not co-vary over animals in relation to the animals’ choice preferences (Fig 7B,C). This finding suggests that drift rate was driven by the animals’ preferences for the 16% liquid sucrose reward, and that the amount of information needed to make a choice or the time taken for sensorimotor processing was not related to the animal’s preferences.

Two other behavioral measures were also evaluated. One was the difference in median latency for trials with low and high-value stimuli. This difference is a proxy for the effect of reward value on the rats’ response latencies. The other was the difference between median latencies for responses to the high-value stimulus on dual and single-offer trials. This difference is a proxy for the effect of choice on the response latencies. These measures did not have large pairwise correlations with any of the DDM parameters. The only significant correlations were between the proxy for choice and the threshold parameter (r=0.5237, df=69, p<0.001, CI95%: 0.33–0.67) and the proxy for value and the drift rate parameter (r=0.3715, df=69, p<0.002, CI95%: 0.15–0.56). These positive relationships suggests that threshold was higher in rats that were slower to respond to the high-value stimuli on dual-offer trials and drift rate was higher in rats that were slower to respond on dual-offer trials compared to single-offer trials.

### Relationships between parameters from the ExGauss and drift diffusion models

Another strong pairwise correlations was found between the Mu parameter (Gaussian) from the ExGauss models and the non-decision time parameter from the DDMs (r=0.6890, df=69, p<0.01, CI95%: 0.54–0.79) (Fig 8A). Interestingly, the other two parameters from the DDMs did not covary with the Mu parameter. Even more dramatic correlations were observed between the Tau parameter (exponential) from the ExGauss models and all three parameters from the DDMs (Fig 8B). Threshold was strongly positively related to Tau (0.8330, df=69, p<<<0.001, CI95%: 0.74–0.89), and therefore threshold was highest in animals that showed the largest exponential variability in their response times. Drift rate and non-decision time showed somewhat weaker negative relations to Tau (Drift Rate: −0.5606, df=69, p<0.001, CI95%: −0.70 to −0.38; NDT: −0.4848, df=69, p<0.001, −0.65 to −0.28), meaning that drift rates and non-decision times were lowest in animals with high levels of exponential variability.

**Figure 8:**
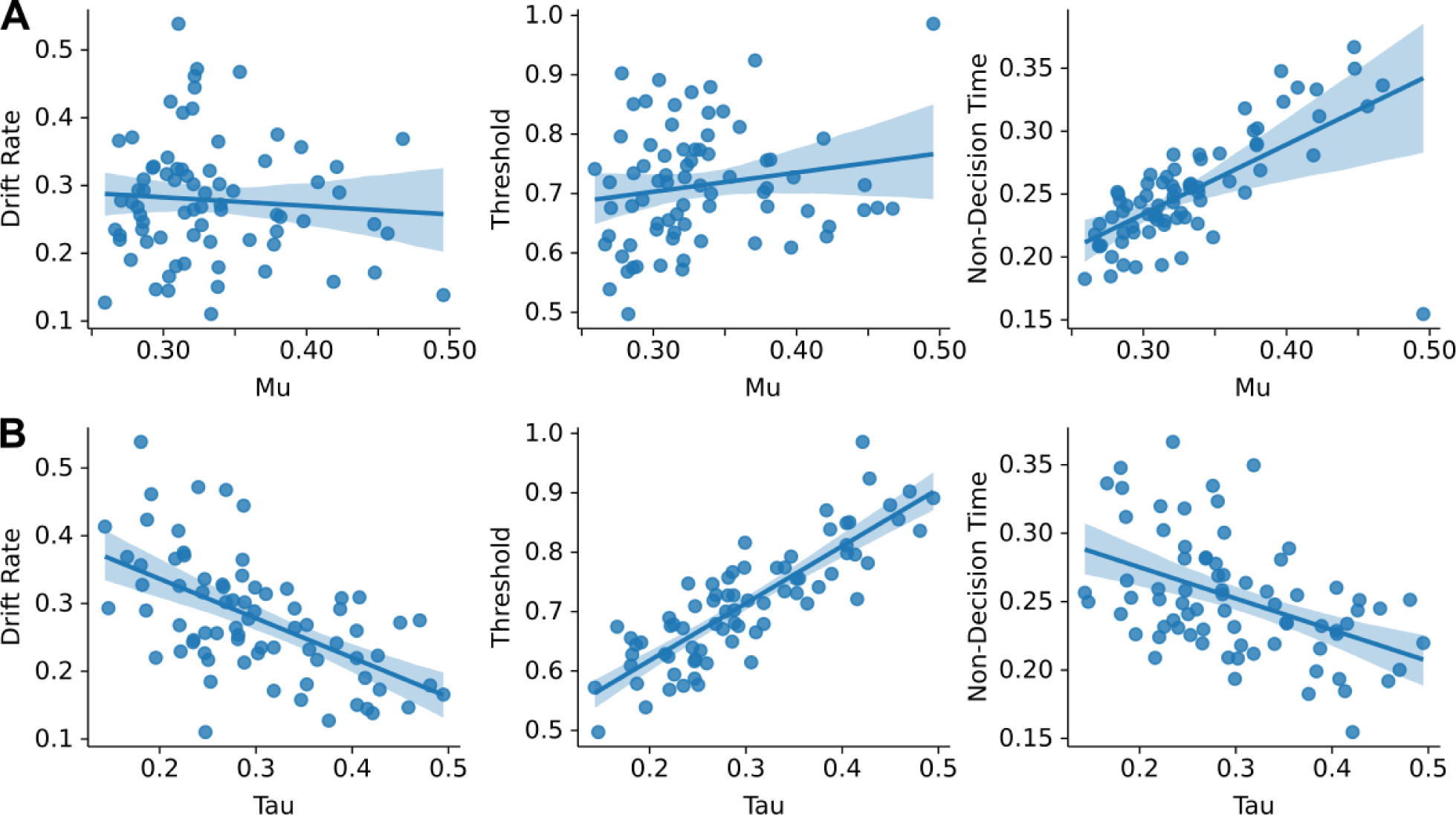
ExGauss parameters had distinct relations to the DDM parameters. A) The Mu parameter from ExGauss modeling was showed a positive relationship non-decision time but not the other DDM parameters. B) The Tau parameter from ExGauss modeling showed strong relationships to all three DDM parameters across animals and sessions.

## DISCUSSION

We studied the behavioral dynamics that underlie initial acquisition of a simple decision-making task. We used a unique training protocol. First, we trained the values of the task stimuli (single-offer trials). Then, we tested animals with choices between pairs of stimuli (dual offers on one-third of trials). The rats took longer to make decisions when faced with these dual offers. With experience, they improved at making choices more quickly. However, the rats still took longer on dual-offer trials compared to single-offer trials. Analysis using drift diffusion models found that less information was required to trigger a choice during the early learning phase. These results do not support the conclusions of Kacelnik and colleagues from their studies of decision making in European starlings (Kacelnik et al., 2011) and several more recent studies done in rats that used a similar training method (Ojeda et al., 2018; Ajuwon et al., 2023). Instead, our results follow what is expected from the Hick-Hyman Law (Hick, 1952; Hyman, 1953), which states that as options increase, response latencies increase, as well as classic studies that motivated the development of drift diffusion models (Ratcliff, 1978). Our findings suggest that brain systems involved in the decision threshold will likely show learning-related changes in neural activity during early choice learning. Neural recording studies are needed to investigate this possibility.

### First instance of choice impacts behavior

During the first session with dual-offer trials, rats showed a preference for high-value cues, indicating they were able to discriminate between cue values when given a choice between the familiar cues in a new context (Fig 2A). Additionally, they showed increased response latencies for dual-offer compared to single-offer trials for both high and low-value trials (Fig 2B). This result suggests rats deliberate (i.e., show slowing on dual-offer trials compared to single-offer trials) and likely engage in some comparative process when deciding between options of known value. Our findings are quite different from the studies by the Kacelnik group that motivated our experiment (Fig 6 in Shapiro et al., 2008; Fig 2 in Ojeda et al., 2018; Fig 4 in Ajuwon et al., 2023). Their studies reported much longer overall response latencies (on the order of seconds). It is possible that the longer latencies reflected a lack of speeded performance and therefore their studies would not reveal an effect of deliberation that is on the order of milliseconds. Our rats responded with median latencies between 520 and 625 ms and showed slowing of ∼100 ms on dual-offer trials with responses to high-value stimuli (Fig 2). An increase in response latency of 100 ms on dual-offer trials added 20% more time to make choices, an effect that is not trivial.

Given the magnitude of difference for latencies to single and dual offers in the first test session, we wanted to assess the stability of that difference and if/how decision making might change with repeated experience. We found there was no significant change in rate of high-value choices on dual-offer trials over five sessions (Fig 3C). This finding suggests the rats were not re-learning the value of each cue when they were presented simultaneously. Their preferences were consistent with the Matching Law (Herrnstein, 1961): The four-fold difference in sucrose concentration used in our study predicts a 75% high-value preference (Graft et al., 1977). We found that over five sessions with dual-offer trials, median response latencies on dual-offer trials reduced, suggesting that the rats learned to choose more quickly (Fig 3A,B). To determine if the effect of deliberation on response latencies we observed in the first session was sensitive to experience, we calculated the difference between single-offer and dual-offer median response latencies per session. We found that the difference was most robust in the first test session and while the magnitude of the difference decreased, we found the effect of deliberation persisted over the five sessions for both reward values (Fig 3D).

Previous studies of visual decision making in rodents have not commonly used the two-stage design (value learning followed by choice learning), as used in the present study. Some of these studies trained animals to detect single stimuli and to report the identity of the stimulus by responding to the left or right, i.e. object-place learning (Zoccolan et al., 2009; Kurylo et al., 2020; Masis et al., 2023). Others trained rodents to make lateralized movements towards the chosen stimulus and trained discrimination between stimuli from the start of initial training (Clark et al., 2011; Reinagel, 2013; Broschard et al., 2019; Broschard et al., 2021; Kurylo et al., 2020; Masis et al., 2023). The only published study that we found that trained rodents with single stimulus presentations before choice learning was Busse et al. (2011). However, that study did not report results on how early task learning affected response latencies.

The changes in deliberation over early choice learning reported in the present study are interesting in the context of many neuroscience studies of decision making that used extensively overtrained animals (see Carandini and Churchland, 2013 for review). If a given brain area is only involved in the acquisition of choice learning, it is possible that it would not show major changes in neural activity once the animals become experienced at performing the decision-making task. As an example, Katz et al., (2016) reported a lack of decision-related firing in area MT in monkeys performing a motion discrimination task. However, if this cortical area was inactivated, the monkeys showed robust impairments in task performance. Area MT is well established as containing neurons that track visual motion (e.g., Anderson, 1997). It is possible that neurons in that cortical area were dramatically altered during the acquisition of the motion discrimination task, and become less engaged after extensive overtraining.

### Choice learning reduced exponential variability and the decision threshold

Two computational approaches were used to quantify how early choice learning affected the animals’ response time and decision processes. Given the distributions of response latencies deviated from the Gaussian distribution (Fig 2C), we implemented a mixture model to estimate the peaks and long tails of the response latency distributions (ExGauss modeling; Heathcote et al., 1991). For high-value trials, there was no difference in the Mu parameter, representing the peak in the response latency distribution (Fig 5A). By contrast, low-value trials showed increased peak latencies for dual-offer trials compared to single-offer trials, over the first four sessions with choice learning (Fig 5B). This finding is suggestive of an overall slowing of when rats chose the low-value stimulus.

The other main ExGauss parameter Tau represents the variability in the tail of the distribution, which might be due to decision variability, aka “noise” in the decision process (Hohle, 1965). Tau was consistently elevated on high-value dual-offer trials compared to high-value single-offer trials throughout the period of choice learning (Fig 5C). Exponential variability decreased over the course of the five choice learning sessions, but the difference between trial types remained significant throughout, suggesting decision variability is elevated on high-value choice trials regardless of the stage of learning. However, in the case of the low-value stimulus, there was no difference for single and dual trial fits of the Tau parameter and no change over the sessions (Fig 5D), so decision variability seemed to only affect high-value choices. Taken together, the two main parameters of the ExGauss models dissociated the higher and lower value trials, a finding that suggests a fundamental difference in how stimuli with different reward values are processed by rats.

To gain insights about how decision-making strategies might change over the course of sessions with dual-offer trials, we fit drift diffusion models to our data (Fig 6). We used two established packages for fitting DDMs, HDDM (Wiecki et al, 2013) and PyDDM (Shinn et al, 2020). We found that the threshold, or boundary separation, was the only parameter that changed over the course of the five sessions in both types of DDM models. This parameter reflects the amount of evidence required to make a choice (Ratcliff, 2001). Our findings suggests that the rats came to require less evidence to respond with repeated experience in making choices, without any changes in drift rate (the rate of accumulation of evidence) or non-decision time (sensorimotor integration not related to the decision process). The finding that drift rate did not change is not surprising given that preference for the high-value stimulus was stable throughout early choice learning, and that there was a strong correlation between drift rate and stimulus preference (Fig 7A) and not the other parameters of the DDM models (Fig 7B,C).

Alternatively, threshold has been associated with caution in performing challenging tasks under time pressure (Forstmann et al., 2008), a process that would depend on inhibitory control. Several recent studies have implicated inhibitory processing in decision making, specifically with regard to the maintenance of the decision threshold (e.g., MacDonald et al., 2017; Roach et al., 2023). By this interpretation, in the present study choice learning might have led to reduced inhibitory control over action. The reduction in inhibitory control would lead to a generalized speeding of performance.

While there was an overall decrease in median response latency over the period of training (Figs 2–3), the results from ExGauss modeling (Fig 5) do not support a role of inhibitory control in choice learning. Specifically, for trials with high-value stimuli, we observed increased exponential, but not Gaussian, variability compared to single-offer trials with high-value stimuli. By contrast, we observed increased Gaussian, but not exponential, variability for trials with choices of low-value stimuli compared to single offers of those stimuli. It is not easy to understand how a common process such as inhibitory control would lead to this dissociation in the Gaussian and exponential components of the latency distributions.

By contrast, the dissociation of effects over the two components of the ExGauss models follows simply from the view that early choice learning reduced the decision threshold. Less evidence would be needed to trigger choices. The rats had already learned to rapidly detect and respond to the high-value stimulus. They did so faster compared to when they responded to the low-value stimulus. The difference between trials with single and dual offers with the high-value stimulus would thus be due to variability arising from the comparison between the stimulus on dual-offer trials. This competition was reflected in the measure of exponential variability captured by the ExGauss models, which was strongly correlated with threshold from the DDMs (Fig 8B).

### Value or visual information?

One limitation of this study was the use of feature matched stimuli and rewards. A high luminance (8 LEDs) cue was paired with a high concentration sucrose reward (16% wt/vol), and low luminance (1 LED) paired with low concentration (4% wt/vol). While this luminance/sucrose pairing is unlikely to impact our main finding about response latencies in choice learning changing over experience, it does bring up the question of whether rats are responding in a perceptual (cue) or value (sucrose) based way, and perhaps why the rats only show a ∼72% preference rate for the high-value stimulus. While we cannot necessarily untangle value cleanly from perception in what is driving response time differences for high and low with our present dataset, unpublished data from our lab suggests a combination effect. In a small cohort of rats (N=6), rats were trained with low-value sucrose mapped to the high luminance and vice versa, the opposite relationship here. We found that rats choose the low luminance, high-value option ∼60% of the time, significantly greater than the low-value, high luminance stimulus (t(29)=9.44, p<0.001), suggesting they are primarily following the value rather than the luminance of each cue.

## METHODS

### Subjects

Thirty male Long–Evans rats (300–450 g, Charles-River, Envigo) were individually housed and kept on a 12 h light dark cycle with lights on at 7:00 A.M. Rats were given several days to acclimate to the facilities, during which they were handled and allowed free access to food and water. During training and testing, animals were on regulated access to food to maintain their body weights at approximately 90% of their free-access weights. All animal procedures were approved by the American University Institutional Animal Care and Use Committee.

Only male rats were used in this study. The behavioral data were collected prior to the implementation of the NIH policy that studies use equal numbers of male and female animals in research projects. The experiments were supported by non-NIH sources of funding. Support from an NIH grant was later in the project when the data analysis was carried out and the manuscript was written.

### Behavioral Apparatus

All animals were trained in sound-attenuating behavioral boxes (ENV-018MD-EMS: Med Associates). A single horizontally placed spout (5/16” sipper tube: Ancare) was mounted to a lickometer (MedAssociates) on one wall, 6.5 cm from the floor and a single white LED light was placed 4 cm above the spout (henceforth referred to as the spout light). The opposite wall had three 3D-printed nosepoke ports aligned horizontally 5 cm from the floor and 4 cm apart, with the IR beam break sensors on the external side of the wall (Adafruit).

Three Pure Green 1.2” 8×8 LED matrices (Adafruit) were used for visual stimulus presentation and were placed 2.5 cm above the center of each nosepoke port, outside the box (see Swanson, et al., 2021 for details about these matrices). Data collection and behavioral devices, including the Arduino that interfaced with the LED matrices, were controlled using custom-written code for the MedPC system, version IV (Med Associates).

### Training Procedure

See Table 1 for a summary of the training procedure and criterion to advance. Animals were initially exposed to 16% sucrose in their homecage to encourage consumption and reduce novelty to the reward during operant training. Rats were then introduced to the operant chambers and trained to lick at a reward spout for 16% wt/vol liquid sucrose in the presence of a spout light and 0.2-sec 4.5kHz SonAlert tone (Mallory SC628HPR). One rat was dropped at this point in the training for lack of interest in consuming sucrose. Over the next several sessions, animals were hand-shaped to respond to dynamic visual stimuli over lateralized nosepoke ports to gain access to a 50µL bolus of liquid sucrose reward at the spout on the opposite side of the chamber. A correct nosepoke (responding at the illuminated port) was indicated by the tone and spout light illumination. These trials comprise the ‘nosepoke responding’ phase of training (Fig 1A). Two rats were dropped from the protocol at this stage of training for not completing >120 trials in a 60 minute session within 5 sessions.

**Table 1:** Training paradigm and criterion to advance rats to the next stage of testing.

| Stage | Name | Goal | Stimulus + Reward | Criterion |
| --- | --- | --- | --- | --- |
| 1 | Continuous Access | Pair spout light and tone with reward | No stimulus; 50 μL 16% liquid sucrose bolus | >500 licks per session |
| 2 | Nosepoke Training - Left or Right Side Only | Rats demonstrate nosepoke responding to operant ports following high luminance stimulus | 8 LEDs over the left or right operant port. Trials only appear on one side port with alternative port blocked, alternate per session. Correct response triggers spout light and tone, indicating availability of 16% sucrose bolus. | >120 trials/session; 3 session minimum, 6 session maximum |
| 3 | Random Cue Presentation | Rats demonstrate they follow the visual cue presentations, not the spatial location. | 8 LEDs presented at random over the L/R operant port. Correct response triggers spout light and tone indicating availability of 16% sucrose reward. Error response to the non-illuminated operant port triggers the spout light requiring response at lick spout, but no reward delivered. | >200 trials/session; <10% error trials; 3 sessions minimum, 6 sessions maximum. |
| 4 | Center Cue Training | Encourage rats to initiate trials by responding to the center 'start' cue. | Start stimulus presented at the center, followed by random presentation of high luminance stimulus over the L/R operant port. Correct and error responses remain the same. | >200 trials/session; <10% error trials; 3 sessions minimum, 6 sessions maximum. |
| 5 | single-offer Only Sessions | Rats are introduced to the low luminance cue and associate it with low-value reward while continuing the same paradigm. | Start cue presented at the center, followed by random presentation of either the high luminance or low luminance cue over the L/R operant port. Response triggers spout light and tone, indicating availability of 16% or 4% sucrose reward, paired to the luminance. Error responses remain the same. | 10 sessions (+/- 2); established response time difference for the high and low response latencies. |
| 6 | Single and Dual Offer Sessions | Investigate whether choice latencies differ for single and dual offer trials for high and low-value trials. | Start cue presented at the center, followed by random presentation of either the high luminance or low luminance cue over the L/R operant port for 2/3 trials. For 1/3 trials, both H/L stimuli are presented over L/R operant ports. Correct and error responses remain the same. | 1 session to 5 sessions of testing |

Rats that reached these criteria were advanced to the next stage of training where a trial initiation cue (4×4 square of illuminated LEDs) was introduced over the central nosepoke. After rats entered the central port, a single stimulus of either high (8 illuminated LEDs) or low luminance (2 illuminated LEDs) was presented without delay. These trials comprise the “value learning” phase of training (Fig 1A). These stimulus presentations were randomized by side and luminance intensity, and importantly were only offered one stimulus at a time (single-offer trial). Correct responses at the port below the high luminance stimulus yielded access to a 16% wt/vol sucrose bolus at the reward spout, while responses at the port below the low luminance cue response yielded access to a 4% wt/vol sucrose bolus. The tone and light were once again indicative of access to these sucrose rewards. Incorrect responses at this stage in the task were indicated only by the presence of the spout light (no tone) and rats were required to make contact with the reward spout before initiating the next trial. Subjects were self-paced throughout the period of training and testing.

When subjects performed >200 trials per 60 minute session during this training phase with fewer than 10% errors, they were moved to the testing phase, “choice learning”, where they were introduced to dual-offer trials (Fig 1A). Two rats were dropped at this stage of training for aberrant strategies (circling all three nosepoke ports), so a total of 25 rats moved on to the testing phase. Ten of these rats were tested in a single session with choice learning. The rest of the rats (N=15) were tested over five choice sessions, with sessions with only single-offer stimuli interleaved over days.

Two types of trials were included in the choice learning sessions. On single-offer trials, animals were presented with a single stimulus randomized by side and luminance intensity. single-offer trials comprised 2/3 of total trials per testing session. On dual-offer trials, which comprised 1/3 of total trials, animals were presented with both the high and low luminance stimuli. The presentation of the brighter stimulus was randomized by side to prevent a spatial strategy in responding on these dual-offer trials.

Response latency was measured as the time elapsed between rats entering the central port to their entrance to their chosen side port after the onset of the visual stimuli. On single-offer trials, we defined errors as entering the non-illuminated port. On dual-offer trials, we determined the choice percentage to assess the preference subjects had for high- and low-value sucrose. See Fig 1 for representation of the training procedure, task, and stimulus sets.

### Data Analysis - software and statistics

Behavioral data were saved in standard MedPC data files (MedAssociates) and were analyzed using custom-written code in Python and [R](https://www.r-project.org/). Analyses were run as Jupyter notebooks under the [Anaconda](https://www.anaconda.com/) distribution.

Statistical testing was performed with R and the [scipy](https://scipy.org/), [pingouin](https://pingouin-stats.org/build/html/index.html), and [DABEST](https://acclab.github.io/DABEST-python/) packages for Python. Repeated measures ANOVA (with the error term due to subject) were used to compare behavioral data measure estimates (median latency, high choice percentage, ExGauss parameters, and DDM parameters from the PyDDM package—see below) across trial type (single or dual offer), value (high or low), and/or session number (for rats tested over 5 sessions). For significant rmANOVAs, the error term was removed and Tukey’s post hoc tests were performed on significant interaction terms for multiple comparisons. Descriptive statistics are reported as mean ± SEM, unless noted otherwise. Two-sided, paired permutation tests were used to compare single and dual offers within each session (Ho et al., 2019) with Bonferroni corrected p-values. 5000 bootstrap samples were taken; the confidence interval was bias-corrected and accelerated. The p-values reported are the likelihood of observing the effect size if the null hypothesis of zero difference is true. For each p-value, 5000 reshuffles of the control and test labels were performed. Results are displayed as summaries with individual points produced by [Matplotlib](https://matplotlib.org/) and [Seaborn](https://seaborn.pydata.org/). Within session effects of trial time and response latency were analyzed with Huber regression from the scikit-learn package for Python. Relations among the parameters from ExGauss modeling and PyDDM were examined using the regplot function from the Seaborn package. Results were quantified using pairwise correlation (Spearman’s rank) using the pairwise_corr function from the pingouin package. Repeated-measures correlation, for the effect of training sessions, was quantified using the rm_corr function from the pingouin package.

### Data Analysis - behavior

Response latency was defined as the time elapsed from the initiation of a trial to the nosepoke response in the left or right port. Response latencies greater than 3 seconds were screened out of the data to exclude trials where rats were disengaged from the task. Median response latency was calculated per rat. Choice percentage reflects the rate at which rats responded to the high luminance target on dual-offer trials. This was calculated by dividing the number of high luminance trials by the total number of dual-offer trials per rat. Errors on single-offer trials occurred when rats produced a nosepoke response in a non-illuminated port. Error percentages were calculated by dividing the number of error trials by the total number of single-offer trials per rat.

### ExGauss modeling of response latencies

Due to the right-skewed nature of response latencies, we wanted to assess how latency distributions change over repeated sessions. Here we used ExGauss model fitting to analyze the distributions. ExGauss is a mixture model of a Gaussian (normal) distribution and exponential distribution, the latter of which captures the extended tail of the response latencies. The [retimes](https://cran.r-project.org/src/contrib/Archive/retimes/) library for R was used for the ExGauss analysis. This package is based on Cousineau et al. (2004). The timefit() function was used to fit an ExGauss model to data for individual rats for each session number (five sessions); number of LEDs (2 or 8); and trial type (single and dual offers). To assess the accuracy of fit of these models to the raw data, especially in cases of a smaller dataset for low-value dual-offer trials, we used the ExGauss() function to generate data from the parameter fits and compared the generated and raw data with a Kolmogorov-Smirnov test was used. Any significant differences (p<0.05) between raw and generated data distributions were further tested with a non-parametric ranksums test. No significant differences were found between the datasets, suggesting the ExGauss parameters were fair to represent the raw distributions. To further validate this toolbox, we also used the [exgfit](https://www.mathworks.com/matlabcentral/fileexchange/70225-exgfit-fit-exgaussian-distribution-to-data) toolbox in MATLAB and found no difference in the parameter fits.

### Drift Diffusion Model fitting using the HDDM package

HDDM models were fit by running version 0.9.6 of the package under Python 3.7. Parameters were from Pedersen et al. (2021): Models were run five times, each with 5000 samples and the first 2500 samples were discarded as burn-in. A fixed bias term of 0.5 was used. Convergence was validated based on the Gelman-Rubin statistic (Gelman and Rubin, 1992). The autocorrelations and distributions of the parameters for each parameter and predictions of the response latency distributions for each animal were visually assessed to further assess convergence.

### Drift Diffusion Model fitting using the PyDDM package

The PyDDM package in Python (Shinn et al., 2020) was used to fit generalized drift diffusion models to these data. PyDDM does not use hierarchical Bayesian modeling. It is an algorithmic approach, based on the Fokker-Planck equation (Shinn et al., 2020). An advantage of using PyDDM was that we could obtain estimates of the decision parameters for each rat, and compare them with other behavioral measures, as reported in Fig 7–9.

We fit an initial model to the response latency distribution from all rats for each session under the method of differential evolution. For our model we fit a total of 3 parameters: drift rate which was linearly dependent upon the log of the measured luminance of the high and low stimuli (drift*log(luminance)), boundary separation, and non-decision time. Our model also accounted for 3 constant parameters: Noise, or the standard deviation of the diffusion process, set to 1.5; Initial Condition set to 0.1 to account for initial bias towards the high-value stimulus; and a general baseline Poisson process lapse rate set to 0.2 specified by the PyDDM package for likelihood fitting. Once this model was tuned to the group data (to identify which parameters may be dependent on task parameters), we fit the model to data from individual rat data from each session to be able to analyze individual shifts in parameters. A repeated measures ANOVA was performed to assess learning effects on the fitted parameters (drift, boundary separation, and non-decision time) and posthoc tests used permutation methods from the estimation statistics package DABEST (Ho et al., 2019).

## DATA SHARING

Data files are available on GitHub: https://github.com/LaubachLab/LearningToChoose

## Acknowledgments

This project was supported by grants from the Klarman Family Foundation and NIH R15DA046375-01A1 and a Faculty Research Support Grant from American University. We thank Drs avid Kearns, Jibran Khokhar, and Elisabeth Murray and Jensen Palmer for helpful comments on this roject and manuscript. We also thank Meaghan Mitchell for assistance in animal care and training.

